# Excessive Postnatal Smooth Muscle Differentiation in a Lung Specific Model of *TBX4*-related Pulmonary Hypertension

**DOI:** 10.1101/2025.03.26.645517

**Authors:** Lea C. Steffes, Kaylie Chiles, Sehar R. Masud, Aleen Rahman, Madeline Dawson, Csaba Galambos, Maya E. Kumar, Ripla Arora

**Affiliations:** Department of Pediatrics, Division of Pulmonary Medicine, Stanford University School of Medicine; Department of Obstetrics, Gynecology, and Reproductive Biology, Michigan State University; Institute for Quantitative Health Science and Engineering, Michigan State University; Departments of Pathology and Pediatrics University of Colorado School of Medicine and Children’s Hospital Colorado Aurora Colorado USA

## Abstract

Heterozygous *TBX4* variants are the second most common genetic cause of pediatric pulmonary hypertension (PH), yet the mechanisms underlying the pathophysiology of TBX4-related lung disease remain poorly understood. We developed a lung mesenchyme-specific *Tbx4* loss of function (*Tbx4cKO*) mouse model that bypasses embryonic lethality to investigate TBX4-related lung disease. Echocardiography of adult *Tbx4cKO* mice demonstrated significant hemodynamic changes consistent with PH. Three-dimensional whole-mount analysis of embryonic day 18.5 lungs revealed reduced lobe volumes and decreased distance between pleural edges and muscularized vessels. In adult *Tbx4cKO* lungs, high-resolution spatial quantitation identified extensive vascular remodeling characterized by significant medial thickening, distal muscularization of small diameter arteries, and extension of muscularized vessels into normally non-muscularized subpleural zones. Contrary to previous reports suggesting vascular simplification with *Tbx4* loss, our comprehensive three-dimensional approach demonstrated an elaborated arterial tree with pathologic muscularization. Additional heterozygous loss of *Tbx5* (*Tbx4cKO;Tbx5he*t) exacerbated vascular phenotypes without worsening hemodynamic parameters. We also documented dysregulated airway smooth muscle patterning and prominent subpleural smooth muscle bands that share molecular features with myofibroblasts and airway smooth muscle cells, echoing pathologic findings in human TBX4 syndrome lung tissue. Collectively, our findings identify TBX4 as a critical suppressor of smooth muscle differentiation across multiple pulmonary compartments. This model recapitulates key features of human TBX4 syndrome and reveals mild developmental underpinnings with subsequent progressive postnatal smooth muscle dysregulation, highlighting a postnatal window during which therapeutic regulation of mesenchymal differentiation may be beneficial.

## INTRODUCTION

Pulmonary hypertension (PH) is a devastating and incurable disease defined by elevated pulmonary arterial pressures of greater than 20mmHg (1). PH is now the leading indication for pediatric lung transplant (2, 3). Advanced PH is hallmarked by arterial changes including increased thickness of the smooth muscle media, muscularization of distal vessels and the formation of occlusive luminal lesions (4). PH is driven by diverse upstream etiologies including genetic mutations, cardiopulmonary comorbidities, systemic inflammatory conditions, drug and toxin exposures and infections, and many cases remain idiopathic (1).

In pediatrics, over the past decade, the increased use of genetic testing has identified novel genetic causes of PH (5). Current research suggests that approximately 40% of children initially diagnosed with idiopathic pulmonary arterial hypertension (PAH) likely have an underlying disease-causing variant (6). These genetic discoveries provide a new lens through which to investigate the molecular mechanisms underlying vascular remodeling in this devastating disease. This deeper understanding could reveal shared and variant-specific pathways, enabling the future development of more effective therapeutics that can either target common disease mechanisms or be precisely tailored to specific genetic variants. Among these genetic causes, heterozygous T-box transcription factor 4 (*TBX4)* variants are particularly prevalent in pediatric-onset PAH, accounting for an estimated 8% of previously idiopathic cases, second only to Bone Morphogenetic Protein Receptor Type 2 variants (6, 7). TBX4 syndrome case series encompassing adolescent and adult-onset PH indicate TBX4-related PH can manifest later in life (8-10). The true prevalence of *TBX4* variants in adult PH populations may be underestimated as genetic testing is not routinely performed.

*Tbx4* encodes a transcription factor expressed in the developing hindlimb, lung, and umbilicus in mice (11, 12). Single-cell analysis has confirmed parallel expression patterns in the human developing hindlimb and lung, though umbilical expression in humans remains to be determined (13, 14). *TBX4* variants were first recognized in Ischiocoxopodopatellar syndrome (15), also known as small patella syndrome, with more recent recognition of lung and pulmonary vascular phenotypes (16). A variable combination of PH, developmental lung disease and hind limb abnormalities are now referred to as TBX4 syndrome (17). In patients with TBX4-related PH, the age of disease onset shows remarkable variability, ranging from fatal neonatal presentations to geriatric-onset disease (8, 9, 18, 19). TBX4 syndrome can manifest with a biphasic clinical picture in which transient neonatal PH and hypoxemia is followed by a ‘honeymoon’ period of clinical wellness before the onset of chronic and progressive PH later in life (19). Since lung biopsies are rarely obtained in infants with resolving neonatal PH, there is limited insight into the extent of developmental vascular abnormalities present during this early phase and whether they persist subclinically during the apparent recovery phase. Classic PAH vascular changes are reported in chronic PH TBX4 patients of all ages, with case reports suggesting a worsening vascular phenotype over time (9, 19, 20). Alongside the vascular phenotype, histologic analysis of TBX4 syndrome lungs shows heterogeneous and patchy developmental lung disease ranging from acinar dysplasia to mild alveolar growth abnormality, as well as ectopic smooth muscle cell (SMC) and even intrapulmonary boney deposition (18-20). It is incompletely known how loss of *TBX4* gene function causes this progressive, wide ranging and distinct lung phenotype (17). A better molecular and cellular understanding of this process could inform clinical interventions to halt disease progression.

In both the human and mouse developing lung, *TBX4*/*Tbx4* is expressed exclusively in the mesenchyme (11, 21-24) and in mouse has been shown to mark the progenitors of vascular SMCs, airway SMCs, fibroblasts, pericytes and chondrocytes (25, 26). No expression is detected in epithelial or endothelial lineages in either human (13) or mouse fetal single cell datasets (24, 27). Postnatally *TBX4* transcription remains mesenchymal, with expression in SMCs, pericytes and fibroblasts (23, 24, 27, 28). This mesenchyme specific localization of *TBX4* in the lung is distinct from the majority of genes associated with PH which are predominantly expressed in the endothelium (5). Loss of TBX4, a quintessential mesenchymal transcription factor, impacts developmental lineages that provide critical signaling and structural support to airways and vasculature both during embryogenesis and postnatally (12). This disruption aligns with the characteristic pathobiology of TBX4-associated lung disease, which manifests as a spectrum of both respiratory and pulmonary vascular abnormalities.

A mouse model of TBX4 syndrome would provide a valuable tool for investigating TBX4-driven lung disease *in vivo*. However, studying the effects of *Tbx4* loss of function on lung pathogenesis is challenging due to the lack of a reported phenotype in heterozygotes and embryonic lethality of homozygous *Tbx4* null mice due to failure of chorioallantoic fusion and formation of a functional placenta (29). Conditional deletion of *Tbx4* using the global *Rosa26*CreER suggests that *Tbx4*-depleted lungs are smaller at mid-gestation in the setting of a failing allantois, but embryonic lethality precluded any postnatal or adult analysis of lung phenotypes (21). To overcome these limitations, we developed a novel TBX4 mouse model by combining a Cre recombinase whose expression is controlled by a lung specific *Tbx4* enhancer (25) with *Tbx4* conditional floxed alleles (29), resulting in removal of *Tbx4* gene function throughout the lung mesenchyme and its derivatives. While this lung-specific deletion differs from human disease, where *Tbx4* is lost globally, it bypasses embryonic lethality and allows investigation of late embryonic, postnatal and adult lung phenotypes. *Tbx5*, a homolog of *Tbx4*, displays overlapping expression with *Tbx4* and is critical for fetal lung development (21). Thus, a *Tbx5* conditional allele (30) was incorporated to examine the effects of loss of an additional T-box homolog on the lung phenotype.

Here, we show that mice with lung mesenchyme specific *Tbx4* deletion develop echocardiographic findings consistent with PH coupled with airway and vascular changes matching those seen in TBX4 syndrome. Through quantitation of artery remodeling from single optical sections, we identified significant medial thickening with *Tbx4* loss of function consistent with changes seen in PAH. Three-dimensional analysis of embryonic day (E) 18.5 mutant lungs revealed that *Tbx4* mutant lungs have smaller lobe volumes and decreased distance between the pleural edge and muscularized arteries and airways. Similar three-dimensional analysis of smooth muscle lineages in adult lungs revealed a progressive phenotype with extensive distal muscularization of an elaborated artery tree, airway muscularization defects, widespread parenchymal myofibroblasts, and extensive subpleural SMC banding—all significantly worsened with the additional loss of a *Tbx5* allele. Each phenotype features excessive smooth muscle deposition, suggesting a central function of lung mesenchymal *Tbx4* expression in appropriately repressing smooth muscle identities. This model of TBX4-related lung disease provides an invaluable tool for studying the mechanisms of PH development in TBX4 syndrome, enabling hypothesis generation and providing a tractable model for *in vivo* hypothesis testing.

## METHODS

### Mice

The *Tbx4*^*LME-Cre*^, *Tbx4* conditional (“floxed”) allele, and *Tbx5* conditional allele mice have been previously described (25, 29, 30). *Tbx4*^*LME-Cre*^; *Tbx4*^*flox/flox*^ (hereafter referred to as *Tbx4cKO*) mice were made by mating *Tbx4*^*LME-Cre*^ males with conditional allele *Tbx4*^*flox/flox*^ females and subsequently crossing *Tbx4*^*LME-Cre*^; *Tbx4*^*flox/+*^ males with *Tbx4*^*flox/flox*^ females to reach homozygosity of *Tbx4* conditional alleles. Animals for analysis were generated by crossing *Tbx4cKO* with *Tbx4*^*flox/flox*^, and Cre negative littermates served as controls. Similarly, *Tbx4*^*LME-Cre*^; *Tbx4*^*flox/flox*^; *Tbx5*^*flox/+*^ (hereafter referred to as *Tbx4cKO;Tbx5het*) mice were made by mating *Tbx4cKO* males with *Tbx4*^*flox/flox*^; *Tbx5*^*flox/flox*^ females, resulting in homozygous *Tbx4* conditional alleles and heterozygous *Tbx5* conditional and wild type alleles. Gestational age was determined by observation of a vaginal plug (designated as day 0.5). All animal maintenance and procedures were performed in accordance with the Institutional Animal Care and Use Committee guidelines at Michigan State University.

### Genotyping & Primers

Excision of *Tbx4*^*flox*^ and *Tbx5*^*flox*^ was analyzed by polymerase chain reaction. Embryonic lungs were dissected at E18.5, and hearts and tracheas were removed. Embryonic lung tissue was lysed in lysis buffer (3% proteinase K) overnight at 55°C, followed by enzyme inactivation for 10 minutes at 95°C. Lysed lung tissue was genotyped for Cre and *Tbx4* as previously described (21). The full list of PCR primers used is listed in Supplemental Figure 1C. The conditional allele was effectively excised in Cre-positive lungs, as indicated by the marked decrease in intensity of the conditional band and presence of the null band (Fig. S1A).

**Figure 1.**
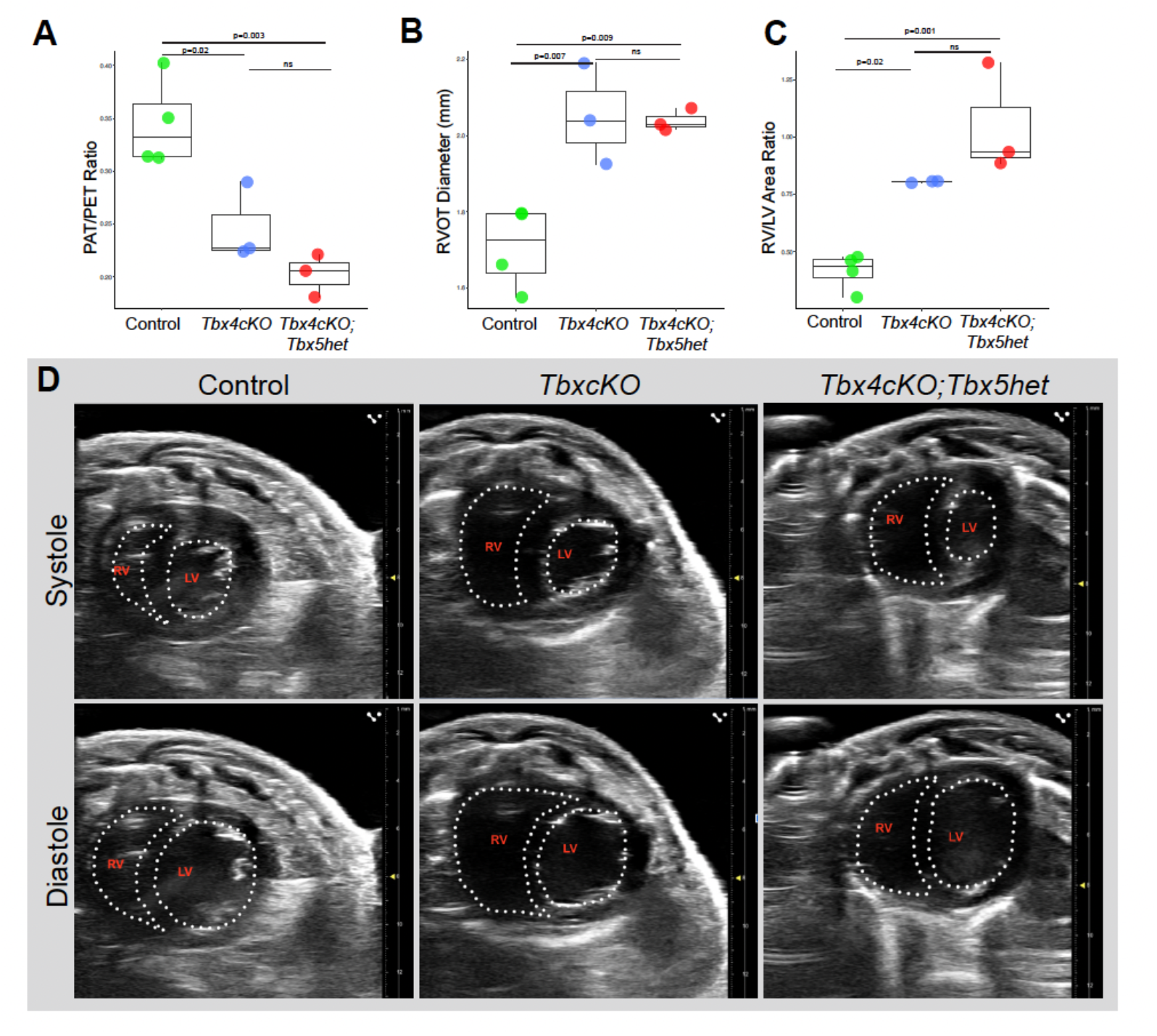
Echocardiographic assessment of adult mice reveals PH and right ventricular dilation in *Tbx4cKO* and *Tbx4cKO;Tbx5het*. **A**, PAT/PET is significantly reduced in *Tbx4cKO* and *Tbx4cKO;Tbx5het* compared to control. Increased right ventricular outflow tract dilation (**B**) and RV/LV area ratio (**C**) in *Tbx4cKO* and *Tbx4cKO;Tbx5het* compared to control. **D**, Representative images from systole and diastole phases of the cardiac cycle depicting the dilated right ventricles in *Tbx4cKO* and *Tbx4cKO;Tbx5het* mice, with flattening of the interventricular septum. Dotted lines denote the ventricular lumen.

### Hemodynamic Assessment by Echocardiography

Cardiac ultrasound imaging (echocardiography) was performed on adult mice. Mice were anesthetized using 4% isoflurane in oxygen at 0.8-2 L/min in a chamber, and anesthesia was maintained using 2% isoflurane in oxygen at 0.8 L/min delivered through a nose cone for the duration of imaging. Animals were kept on a heated platform while under anesthesia and heart rate was monitored using electrocardiogram leads. Fur was removed from the thorax using electric clippers immediately prior to imaging. A VevoF2 (Fujifilm VisualSonics) high-frequency ultrasound with a UHF57x (25-57 MHz) transducer was used to image the hearts and great vessels in short axis and long axis parasternal views. The Doppler function was used to assess cardiac blood flow (31).

### Embryonic tissue collection

Embryonic lungs were collected from timed matings at E18.5. Pregnant mice were euthanized at E18.5 using carbon dioxide asphyxiation and cervical dislocation. Embryos were dissected from the uterus with yolk sac intact and immediately placed on ice. Lungs were dissected from embryos in ice cold 0.1% bovine serum albumin in PBS. Lung lobes were separated, fixed in 1:4 DMSO:methanol, and stored at -20°C overnight.

### Adult tissue collection, perfusion, inflation, fixation and storage

Adult lungs were collected from age-matched littermate *Tbx4cKO* or *Tbx4cKO;Tbx5het* mice and controls between 4-9 months of age. Mice were euthanized using carbon dioxide asphyxiation and lungs were perfused using ice cold PBS. Lungs were inflated using 2% low melting point agarose in PBS, placed on ice and lung lobes were separated and fixed individually. Caudal lobes were fixed in 4% PFA followed by serial dehydration in 3:1, 1:1, and 1:3 solutions of PBS:methanol, then placed in 100% methanol and stored at -20°C until ready for staining. Left lungs were fixed in 4% PFA followed by serial dehydration in 10%, 20%, and 30% solutions of sucrose in PBS, then embedded in OCT and stored at -80°C until ready for staining.

### Histology and immunohistochemistry

Cryosections were prepared and stained following standard protocols (32). Briefly, 4% PFA fixed lobes were cryopreserved in 30% sucrose in PBS, embedded in OCT (Sakura, 4583) and 20μm sections were cut using a Leica CM3050S cryostat. Cut sections were stored at -80°C prior to staining. On the day of staining slides were thawed, washed in PBS + 0.1% Tween-20, blocked at least 30 min in preblock consisting of 0.3% Triton X-100, 5% serum and 1.5% BSA, and then incubated overnight in primary antibody solution diluted in preblock at room temperature. The next day slides were washed, incubated 45 minutes in secondary antibody solution, nuclei were stained with DAPI (1:1000 dilution of 10 mg/ml stock; Invitrogen, D1306), and slides were mounted with Prolong Gold antifade mounting media (ThermoFisher, P36930). For visualization of elastin fibers by staining with fluorescent hydrazide dye (33), stock solutions were prepared by dissolving 1mg hydrazide-A633 (Invitrogen, A30634) in 2ml diH_2_O, and were used at a 1:500 dilution in the primary or secondary antibody mix.

Primary antibodies used: mouse anti αSMA-Cy3 (Sigma, C6198; 1:200), mouse anti αSMA-FITC (Sigma, F3777; 1:200), rat anti mouse CD31 (BD Pharmingen, 553370; 1:500), hamster anti mouse CD31 (BioRad, MCA1370Z; 1:200). Secondary antibodies used: A647-conjugated goat anti hamster (Invitrogen, A-21451; 1:250), A488 goat anti rat (Invitrogen, A-11006; 1:250).

### Whole mount antibody staining

Whole mount immunofluorescent staining of E18.5 lungs was performed as previously described (34). Embryonic lung lobes fixed in DMSO:methanol were rehydrated in 1:1 methanol:PBT (1% Triton in PBS) for 15 minutes, then washed in PBT for 15 minutes at room temperature. Lung lobes were then incubated in 2% powdered milk in PBT blocking solution for 2 hours. Blocking solution was removed and primary antibodies were added (1:250 antibody dilution in blocking solution). Lung lobes were incubated in primary antibody, mouse anti αSMA-FITC (Sigma, F3777; 1:250), for five nights at 4°C. Lung lobes were then washed in PBT twice for 15 minutes and four times for 45 minutes each at room temperature. Hoechst (B2261, Sigma-Aldrich; 1:250) was diluted in PBT; lung lobes were incubated for three nights at 4°C. Next, lung lobes were washed in PBT twice for 15 minutes and four times for 45 minutes each at room temperature, then dehydrated in 1:1 methanol PBT followed by 100% methanol for 30 minutes. Lung lobes were subsequently bleached in 3% hydrogen peroxide in methanol at 4°C overnight. Lastly, bleached samples were washed in 100% methanol for 30 minutes and cleared overnight in 1:2 benzyl alcohol:benzyl benzoate (BABB).

Adult lung lobes stored in 100% methanol were rehydrated, bleached in 5% hydrogen peroxide in PBS, washed in PBS, incubated with preblock consisting of 0.3% Triton X-100, 5% serum and 1.5% BSA, and then incubated for five nights in primary antibody solution containing αSMA-Cy3 (Sigma, C6198; 1:200) and hydrazide-A633 (Invitrogen, A30634; 1:500 dilution of 0.5mg/ml stock) diluted in preblock at 4°C. Samples were then washed, postfixed briefly in 4% PFA, dehydrated to 100% methanol and cleared in BABB.

### Confocal imaging and quantitation

Embryonic lung samples were imaged using a Leica TCS SP8 X Confocal Laser Scanning Microscope System with white-light laser and 20x BABB immersion objective. The full volume of the lung lobes was imaged using tile scans with z step size of 7μm. Tile scan images were merged using Leica software LASX version 3.5.5. (35) Image analysis for embryonic lung lobes was performed using Imaris v9.2.1 by importing confocal image (.LIF) files into Imaris Surpass mode. Artery and airway SMC metrics were analyzed by creating a surface for smooth muscle and the pleural edge. Caudal lobe volume was analyzed by creating a surface of Hoechst staining and using surface statistics in Imaris.

All immunohistochemistry and RNAscope images were captured on a Zeiss 880 Examiner confocal microscope and minimally processed with Zen software (Carl Zeiss AG).

### Artery morphology assessment from cryosection stains

Media thickness and percent artery diameter for neointima were calculated from single plane confocal images as follows. 20μm cryosections were prepared from lungs of all three genotypes and stained with hydrazide to mark elastic laminae, αSMA to mark SMCs and neointima, CD31 to mark endothelium, and DAPI to mark nuclei. Stained sections were imaged at high resolution by tiled confocal microscopy, and all artery cross-sections with diameters between 20μm and 200μm were identified for measurement in the stitched files. For each artery the following measurements were taken manually using Zen Black (Carl Zeiss AG) software: external vessel diameter (measuring from outside margin of external elastic laminae), media thickness (inside margin of external elastic lamina to outside margin of internal elastic lamina), neointima thickness (inside margin of internal elastic lamina to the outside edge of the endothelium). Two sets of orthogonal measurements were taken for each artery and averaged. Three animals were analyzed for each genotype and for each animal 25-50 arteries were measured.

### Stereoscope imaging and quantitation

For adult whole lobe analysis and detailed analysis of SMC and elastin distribution along peripheral arteries and airways, multi-focus stereoscopic images of the specified branches were collected using a M205 FA fluorescent stereoscope (Leica Microsystems), Orca-Flash 4.0 LT monochrome digital camera (Hamamatsu) and LASX software (Leica Microsystems).

Measurements were made using Fiji, a distribution of ImageJ. Branches that were selected for analysis are ones that are arrayed in a single flat plane, such that measurements from multi-focus images accurately capture the distance to pleura.

### Statistical analysis

Statistical analysis for medial and neointimal thickness, artery and airway distance to the pleura, distal artery diameter and airway SMC ring gapping was performed using Welch’s ANOVA (assuming unequal variances) to assess differences across genotypes, followed by Games-Howell post-hoc tests for pairwise comparisons between groups. For lobe size, mouse weights and hemodynamic parameters statistical analysis was performed using one-way ANOVA across genotypes, followed by Tukey’s HSD post-hoc tests for pairwise comparisons between groups. All statistical analysis and plotting were done in the statistical language R (www.R-project.org) using RStudio (RStudio Inc). Plots were made using the ggplot2 R package (ggplot2.tidyverse.org).

### Acquisition, processing and staining of human tissue

Lung tissues were obtained at autopsy, fixed in 10% neutral buffered formalin and processed, embedded, sectioned and H+E stained following standard protocols (20). αSMA immunostaining was done via automated staining using a Ventana BenchMark XT automated slide stainer (Leica Microsystems).

### RNA *in situ* hybridization

Fluorescent *in situ* hybridization on 20μm cryosections sections was carried out using an RNAscope v2 kit (ACDbio, 323100), following manufacturer instructions. RNA *in situ* hybridization protocol was followed by a 5 min DAPI stain and overnight incubation with αSMA-FITC antibody (Sigma, F3777) diluted 1:200 in preblock solution. Mouse probes used were 483791 (*Cnn1*), 425171 (*Notch3*), 411381 (*Pdgfrb*), 404961 (*Lgr6*), and 448441 (*Hhip*), all from ACDbio.

## Results

### Mice lacking pulmonary *Tbx4* gene function have PH

To assess excision at the *Tbx4* locus using *Tbx4*^*LME*^*-Cre* semi-quantitative PCR-based genotyping of entire embryonic lungs at E18.5 was used. DNA fragments were amplified for *Tbx4* wildtype, floxed (conditional) or null (excised) alleles (21). A marked reduction in the floxed band and a prominent null band indicates efficient excision at the *Tbx4* locus within the pulmonary mesenchyme (Fig. S1A). As endothelial and epithelial lineages do not express *Tbx4*^*LME*^*-Cre*, a band for the floxed allele is expected in both *Tbx4cKO* and control samples. *Tbx4cKO* pups were born in Mendelian ratios, with 50.8% of pups from the cross *Tbx4*^*LME*^*-Cre; Tbx4*^*flox/flox*^ X *Tbx4*^*flox/flox*^ inheriting Cre (n=62/122 Cre-positive pups weaned from 12 litters). Equal proportions of males and females were born (29/62 Cre-positive pups were female and 33/62 pups were male) and *Tbx4cKO* mice had no differences in weight compared to control mice (Fig. S1B).

To further characterize the impact of fetal loss of *Tbx4* and *Tbx5* on adult lung function and the development of PH, we performed echocardiography on adult mice between 4 and 6 months of age. While PH in humans is diagnosed with right heart catheterization, echocardiography is a non-invasive and reliable tool for assessing right ventricular hemodynamics and morphology in mice (36-38). PH metrics collected from mouse echocardiography include the ratio of pulmonary acceleration time to pulmonary ejection time (PAT/PET; a measurement of right ventricular afterload; a decrease in this value is correlated with increased severity of pulmonary hypertension), right ventricular and outflow tract dilation, and leftward deviation of the interventricular septum (36-38). We found that *Tbx4cKO* adult mice exhibited a significant reduction in PAT/PET compared to controls (Fig. 1A; n=3 *Tbx4cKO* and 4 controls; p=0.02 compared to control), a significant increase in right ventricular outflow tract diameter (Fig 1B; p=0.007 compared to control), and a profound increase in the ratio of right ventricular to left ventricular area in the short axis view (Fig 1C; p=0.02). Concurrent loss of one *Tbx5* allele (*Tbx4cKO;Tbx5het)* maintains the *Tbx4cKO* PH phenotype without a statistically significant worsening of hemodynamic parameters (Fig. 1A-C; n=3 *Tbx4cKO;Tbx5het* PAT/PET p=0.003 vs control and p= 0.33 vs *Tbx4cKO*, RVOT outflow tract dilation p=0.009 vs control and p=0.98 vs *Tbx4cKO*, right ventricular to left ventricular area p=0.001 vs control and p=0.15 vs *Tbx4cKO*). Representative short axis echo images by genotype are shown in Figure 1D. No significant changes in cardiac output (CO), stroke volume (SV), ejection fraction (EF) or fractional shortening (left ventricular diameter change during systole) were observed between genotypes, indicating that left ventricular function is not compromised (Fig. S2; n=4 for control and n=3 for *Tbx4cKO* and *Tbx4cKO;Tbx5het*).

### Right caudal lobes are smaller with distal-most artery and airway SMCs found closer to the pleural surface at E18.5

*TBX4* loss of function is associated with lethal developmental lung disorders hallmarked by hypoplastic lungs and varying degrees of acinar developmental failure (18). To identify changes at the end of embryogenesis, *Tbx4cKO* mice were harvested at E18.5, just before birth. Using whole mount immunofluorescence staining, confocal microscopy, and image segmentation, we analyzed right caudal lobe volume and smooth muscle α-actin (αSMA) expression around the distal arterioles and airways in uninflated lungs. Lobe volume was significantly smaller in *Tbx4cKO* mice compared to control (Fig. 2C; average 6.02×10^9^ μm^3^ vs 8.20×10^9^ μm^3^; p = 0.029) and a significant difference in the proximity of artery and airway SMCs to the pleura was observed in *Tbx4cKO* lobes compared to control (Fig. 2 A-B). The shortest distance from the distal-most point of fully coherent arterial SMC coverage to the pleural edge was measured in arteries associated with the terminal bifurcations of airway branches RCd.L1 through RCd.L4 (39). *Tbx4cKO* mice exhibited a significant reduction in muscularized artery-to-pleura distances compared to control (Fig. 2E; p = 0.032), with coherent artery SMC coverage terminating a median distance of 396μm from the pleural edge in control lobes, and 322μm in *Tbx4cKO*. Artery diameter at the point of distal-most coherent artery SMC coverage was measured and no significant change was found in *Tbx4cKO* mice compared to control (Fig. 2D; p = 0.97). In addition to distal artery SMC coverage, we assessed distal airway SMC coverage. Like artery SMCs, distal-most airway SMC coverage was significantly closer to the pleural edge in *Tbx4cKO* compared to control (Fig. 2F; median of 403 μm in controls; 393.5 μm in *Tbx4cKO*; p = 0.044; n=3 controls and n=4 *Tbx4cKOs*).

**Figure 2.**
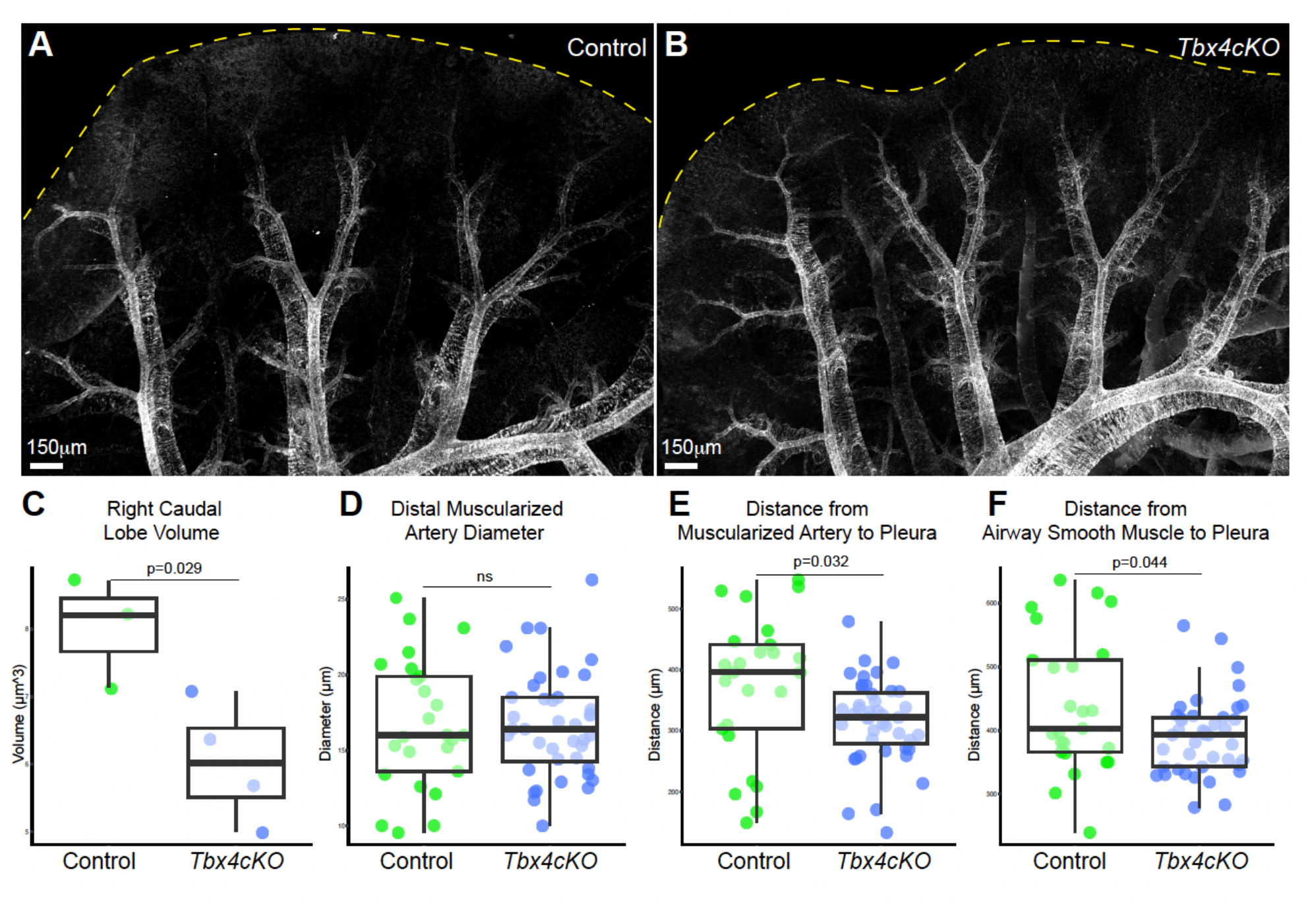
Decreased distance between muscularized airways and arteries and the pleura edge, as well as reduced lobe volume at E18.5 in *Tbx4 KO*. Confocal projections of whole right caudal lobes stained to highlight αSMA (white), cleared, and visualized in three dimensions from control (**A**), Tbx4cKO (**B**), showing αSMA-positive airways and arterioles nearer to the pleural edge in *Tbx4cKO* compared to control at E18.5. **C**, Caudal lobe volume is significantly reduced in *Tbx4cKO* compared to control (p = 0.029). Distal muscularized artery diameter did not differ significantly between genotypes, **D**; however, the distance from distal-most vascular (**E**) and airway (**F**) smooth muscle to the pleural edge was significantly reduced in *Tbx4cKO* compared to control.

### *Tbx4cKO* and *Tbx4cKO;Tbx5het* adult mice have significant medial thickening and variable neointimal formation

Increased muscularization and arterial remodeling is seen in patients with TBX4-related pulmonary vascular disease ((9, 19, 20); Fig 3E&F). Thickening of the smooth muscle media and formation of neointimal lesions, which are cellular accumulations between the intima and internal elastic lamina, are hallmarks of human PAH histology and drive PH hemodynamics (4). To uncover the cause of elevated pulmonary artery pressures observed in adult *Tbx4cKO* we performed a detailed examination of artery anatomy in these animals (Fig 3A-C). Though average artery diameter was comparable between genotypes (56.7μm in control vs. 55.9μm in *Tbx4cKO*), media layers were significantly thicker in *Tbx4cKO* animals, with an increase in mean thickness of 24% compared with controls (Fig. 3D; 3.6μm in *Tbx4cKO* and 2.9μm in control, p= 1.03 × 10^−7^). There is no significant increase in medial thickness in *Tbx4cKO;Tbx5het* compared to *Tbx4cKO* (p=0.11). In one *Tbx4cKO* animal frequent, partially occlusive neointimal lesions were observed (neointima present in 38% of measured arteries; 11 of 29 arteries scored), but neointima was largely absent in all other mutant animals evaluated (neointima present in 0.31% of all remaining arteries; 1 of 318 arteries scored; n=3 for each genotype).

**Figure 3:**
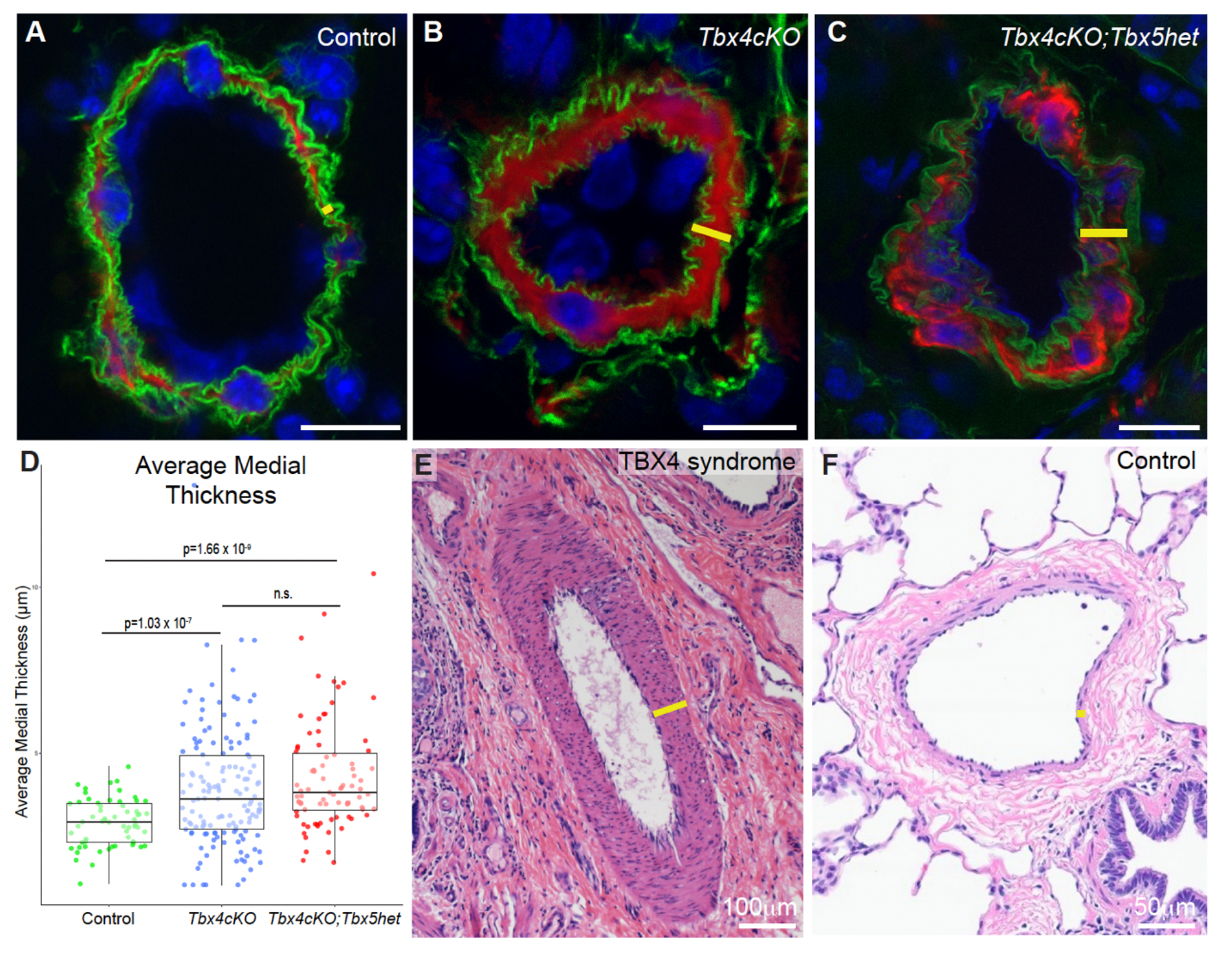
*Tbx4cKO* and *Tbx4cKO;Tbx5het* mice display pulmonary artery muscularization consistent with PH hemodynamics and similar to that seen in TBX4 syndrome. **A**-**C**, Pulmonary artery cross-sections from control, *Tbx4cKO*, and *Tbx4cKO;Tbx5het* mouse lungs stained to highlight media (αSMA, red), internal and external elastic laminae (hydrazide, green), and nuclei (DAPI, blue) show an increase in medial thickness (yellow bars) with loss of *Tbx4* gene function from lung mesenchyme. **D**, Measurement of medial thickness from high resolution confocal images of individual arteries from each genotype confirms medial hypertrophy with loss of *Tbx4*. The additional loss of one copy of *Tbx5* does not lead to significant worsening of the phenotype. **E&F**, Pulmonary arteries of TBX4 syndrome patients display a similar expansion of the medial layer (yellow bars) versus control. A-C, confocal micrographs of cryosections, scale bars 10μm; E&F, H&E stains of paraffin sections.

### Whole mount analysis of *Tbx4cKO* and *Tbx4cKO;Tbx5het* lungs demonstrates excessive vascular SMC coverage and distal muscularization of small arteries

To assess the distribution and extent of muscularized arteries with fine spatial precision whole mount αSMA antibody staining, deep tissue imaging, and detailed quantitation of intact right caudal lobes were performed. A striking feature of the mutant phenotype was the abnormal proximity of fully muscularized arteries to the pleural surface (Fig. 4A-C), paralleling observations in TBX4 syndrome in which muscularized arteries abut the pleura (Fig. 4D-E). As in the embryonic analysis above, to allow quantitative comparison between genotypes we assessed the arteries accompanying the terminal bifurcations of airway branches RCd.L1 through RCd.L4 (39) in all individuals. In control lungs, arteries have a coherent αSMA+ media until a median distance of 511 μm from the pleura, after which only sparse, discontinuous αSMA+ cells are observed. However, in *Tbx4cKO* mice, we found coherent muscularization extended a median distance of 278 μm (p = 6.57 × 10^−9^ vs control) from the pleural edge (Fig. 4F). Many fully muscularized arteries extended into what is normally a non-muscularized vascular zone, with 39.1% of arteries showing coherent smooth muscle media within 160μm of the pleura in *Tbx4cKO* compared to 0 in control. We termed this normally artery muscle-free peripheral region the Subpleural Non-Muscularized Zone (SNZ), demarcated as a dashed line in Figure 4F.

**Figure 4:**
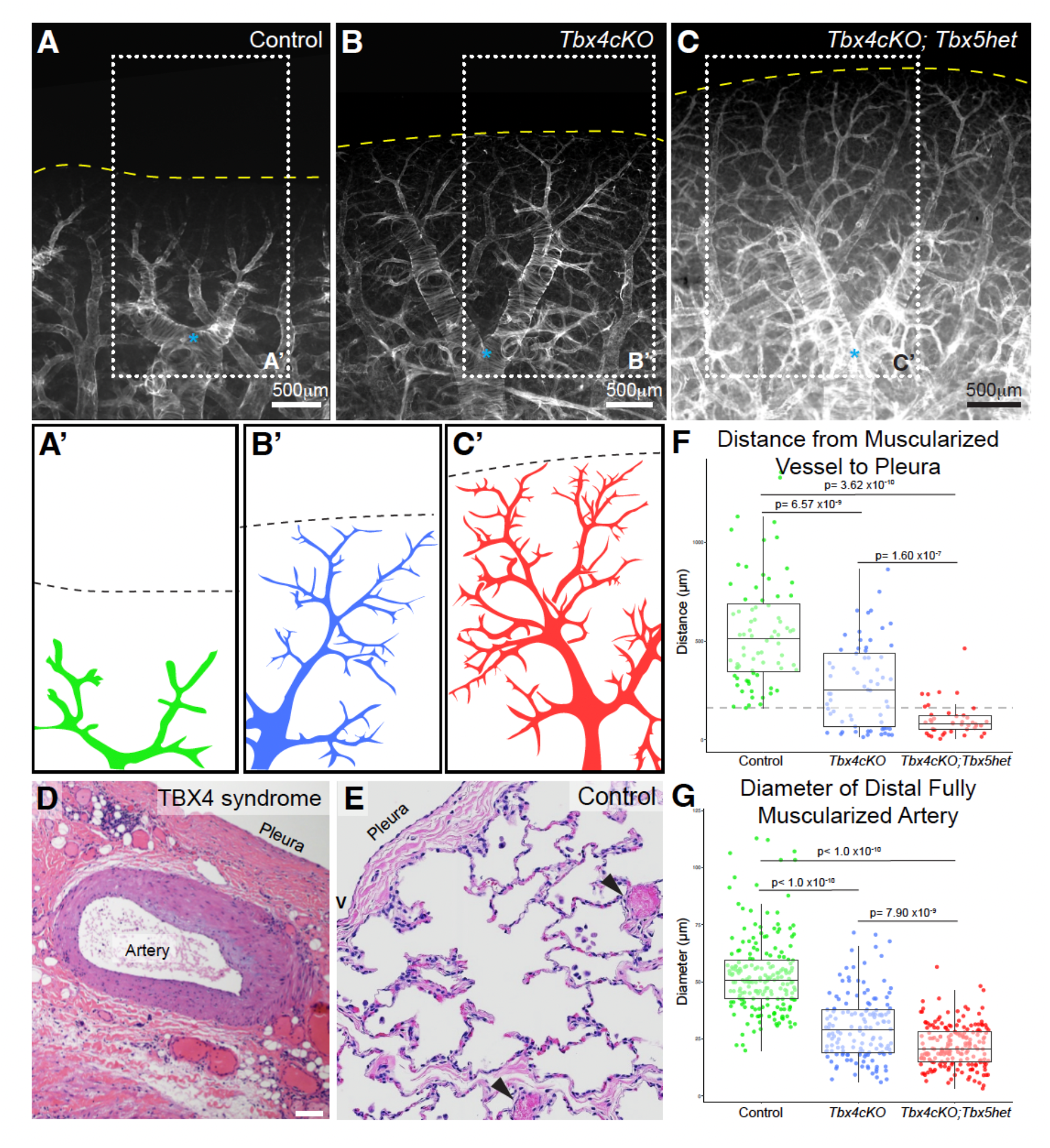
Whole mount analysis reveals increased vascular smooth muscle coverage of peripheral pulmonary artery networks in *Tbx4cKO* and *Tbx4cKO;Tbx5het*. **A**-**C**, Intact right caudal lobes stained to highlight smooth muscle α-actin protein (white), cleared, and visualized in three dimensions from control, *Tbx4cKO*, and *Tbx4cKO;Tbx5het* animals allows visualization of both airway, arterial and venous smooth muscle networks. The same branch (dorsal view of RCd.L3) is shown for each genotype with distal-most airway bifurcation marked with asterisk to aid comparison. Arterial networks for each genotype are schematized in **A’**-**C**’. Highly muscularized arteries are found close to the pleural surface in TBX4 syndrome lungs (**D**) but not in control (**E**). Arrowheads highlight peripheral vessels in E. **F**, In both *Tbx4cKO* and *Tbx4cKO;Tbx5het*, muscularized arteries extend significantly closer to the pleura (yellow dotted lines in A-C; black dotted lines in A’-C’). Gray dotted line in F shows the position of the most peripheral muscularized artery in control lungs. **G**, Consistent with abnormal smooth muscle coverage of slender distal pulmonary arteries, the diameter of the distal-most fully muscularized arteries drops significantly in both *Tbx4cKO* and *Tbx4cKO;Tbx5het* animals. A-C, multifocal projections of stereoscope z-stacks; D&E, H&E staining of paraffin sections.

Distal muscularization, in which smaller diameter arteries have a complete SMC coat, is seen in PAH histology and is thought to be an important driver of increased pulmonary vascular resistance (4). We assessed distal muscularization by measuring the diameter of arteries at the point where the coherent artery SMC coat ends and found that smaller diameter arteries were fully muscularized in *Tbx4cKO* compared with controls (Fig. 4G; median of 30.2 μm in *Tbx4cKO* vs 53.1μm in control; p < 1.0 × 10^−10^).

All artery muscularization parameters were exacerbated in *Tbx4cKO;Tbx5het* with the distance to the pleura of muscularized arteries shrinking to a median of 100μm (p = 1.60 × 10^−7^ vs KO, p= p 3.62 × 10^−10^ vs control), the percent of muscularized vessels within the SNZ increasing to 83.7% (Fig. 4F) and the diameter of muscularized tips narrowing further to 21.9μm (Fig. 4G; p = 7.9 × 10^−9^ vs KO, p < 1.0 × 10^−10^ vs control; control n=7, *Tbx4cKO* n=5, *Tbx4cKO;Tbx5het* n=3). Together, these findings indicate a role for T-box gene function in restricting the distal extent of artery muscularization.

### Excessive artery branching and abnormal patterning of muscularized arteries in *Tbx4cKO*

Arteries of all sizes have elastic laminae which disappear only as they transition to the capillary bed (40). Staining these elastin layers allows us to visualize the arterial tree in whole mount samples independent of the presence of SMCs. To allow direct comparison between control and mutant samples, we quantitated branch number and SMC coat of the same artery (the artery serving the second ventral airway branch emerging from the first lateral branch of the right caudal lobe; RCd.L1.V2; branch number determined by counting all artery branch points visible with either αSMA or elastin staining in the artery tree stemming from the first lateral branch) in control and *Tbx4cKO*. In control, distal arteries are αSMA-negative, with 44% of elastin-marked artery branches lacking a SMC coat (26 elastin-marked branches, 15 αSMA-coated). *Tbx4cKO* shows a more elaborate and muscularized arterial tree with a greater number of elastin-marked branches all of which are invested with a SMC coat. In *Tbx4cKO* the SMC coat extends beyond the visualizable elastic lamina with 47 elastin-marked branches scored but 56 αSMA-covered branches visualized.

### Airway muscularization extends closer to pleura in *Tbx4cKO* and *Tbx4cKO;Tbx5het* compared to control and myofibroblasts are abundant in *Tbx4cKO*

Because bronchioles abutting the pleural edge are observed in TBX4 patients (Fig.5A-B; (19)) we investigated the distribution of SMC-invested conducting airways in relation to the pleura in *Tbx4cKO*. Parallel rings made up of closely spaced bundles of airway SMCs wrap conducting airways (Fig. 5C-D). Moving from proximal to distal, as conducting airways transition to respiratory epithelium, the SMC rings abruptly become sparse and are absent in the most peripheral airways (41). In *Tbx4cKO* animals, airway SMC rings are present closer to the pleura than in control (Fig. 5E; median of 731μm in KO vs 1188μm in control, p= 2.77 × 10^−4^). Additionally, the spacing of SMC rings was altered in *Tbx4cKO*, with wider spacing of rings both proximally (median gaps of 25.6μm in KO vs 19.7μm in control, p= 1.00 × 10^−3^) and distally (median gaps of 80.9μm in KO vs 25.6μm in control, p= 1.04 × 10^−5^) relative to a fixed position in the airway tree, the terminal bifurcation of branches RCd.L1.Cr1, RCd.L1.Cr2 and RCd.L1.Cr3 (39) (Fig. 5F). Measurement of gap distances and the distal extent of airway SMC rings in *Tbx4cKO;Tbx5het* was unreliable due to the large number of αSMA+ myofibroblasts throughout the parenchyma. (Airway to pleura measurements: control n=7, *Tbx4cKO* n=4; SMC gap distance measurements: control n=3, KO n=3). In addition, dispersed C-shaped αSMA-positive cells not present in controls were found throughout the parenchyma of *Tbx4cKO* lungs (Fig. 5G-H) with further expansion in *Tbx4cKO;Tbx5het* mice. These cells were not associated with conducting airways, arteries or veins and are inferred to be myofibroblasts.

**Figure 5:**
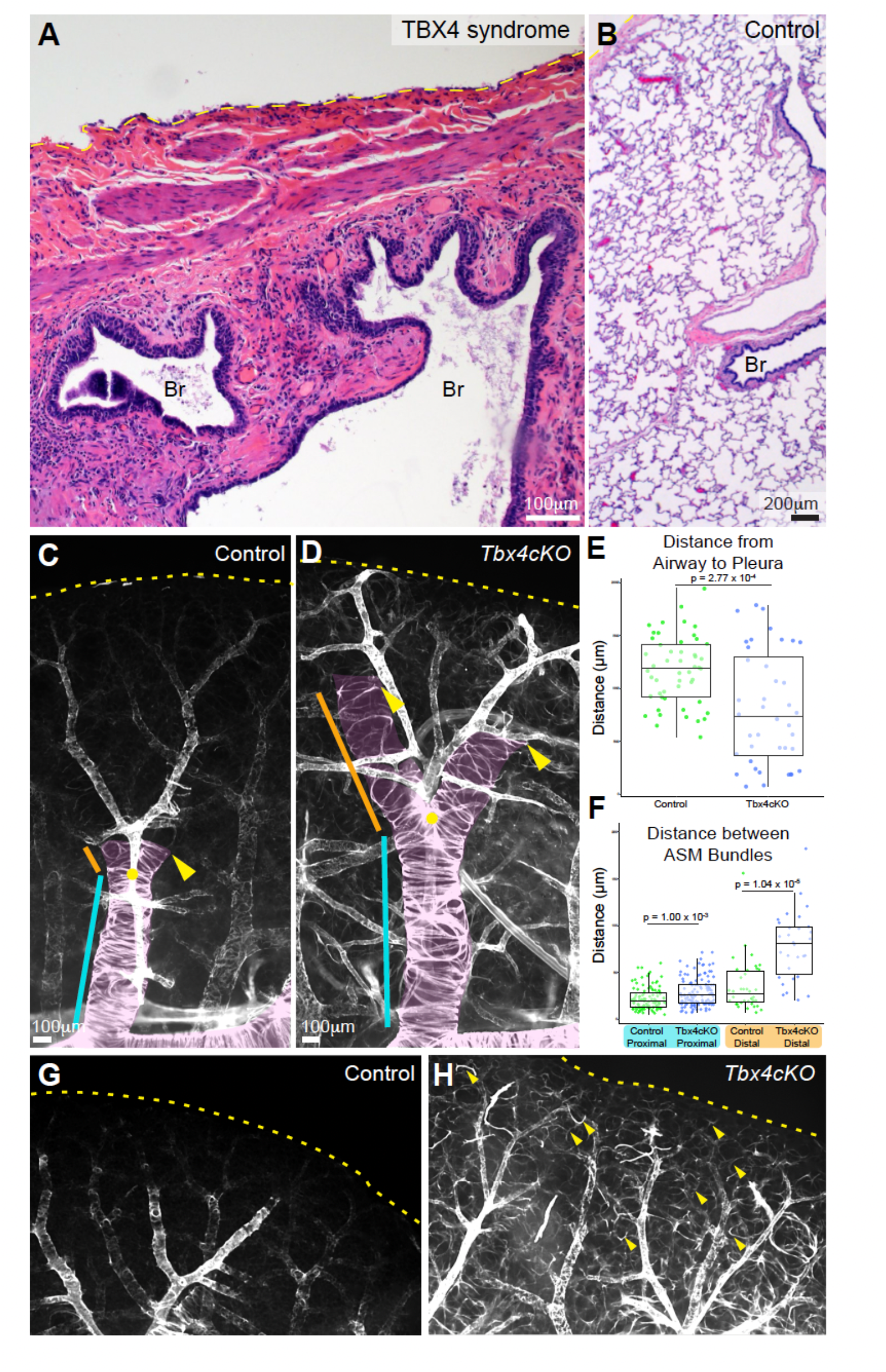
Airway smooth muscle extends closer to the pleura and is mispatterned in *Tbx4* mutant lungs. **A&B**, In TBX4 syndrome smooth muscle-wrapped conducting airways are found in close apposition to the pleura, while in control conducting airways end more proximally. **C&D**, whole mount stains of right caudal lobes highlight αSMA+ cells (white) along airways and vasculature. The same branch (dorsal view of RCd.L1.Cr2) is shown in each genotype, with airway associated smooth muscle indicated with pink overlay and the position of the distal-most complete smooth muscle ring indicated with yellow arrowheads. **E**, The distance between distal-most smooth muscle ring and the pleural margin is significantly shorter in *Tbx4cKO* animals versus control. **F**, The distance between smooth muscle bundles both before (cyan bar in C&D) and after (orange bar in C&D) the final conducting airway bifurcation (yellow dot, C&D) is significantly wider in *Tbx4cKO*. **G&H**, whole mount αSMA (white) staining of right caudal lobe margins shows an abundance of αSMA+ C-shaped myofibroblasts (selected cells highlighted with arrowheads) in parenchyma of *Tbx4cKO* not present in control. Pleural lobe margin indicated with yellow dotted line in A-D, G&H.

### Pleural SMC bands are found with increasing density with T-box allele loss and express markers of myofibroblasts and airway SMCs

In addition to the airway and vascular abnormalities described above, whole mount αSMA antibody staining of intact lobes revealed the presence of αSMA-expressing bands, predominantly running along the ventral face of the pleural surface. These bands were composed of long strands of parallel smooth muscle-like fibers grouped in coherent bundles (Fig. 6E). Short, thin bands were present in control specimens (Fig. 6A). However, in *Tbx4cKO* and *Tbx4cKO;Tbx5het*, we observed a graduated and striking increase in the length, density, and number of these pleural fibers (Fig. 6B-C). Though bands were prominent in all *Tbx4cKO;Tbx5het* lobes examined, in one animal elaborate webs of SMC bands covering the majority of the pleural surface were observed, to the point of obscuring visualization of underlying structures with fluorescence microscopy (Fig. 6C). These bundles wrap around lobe edges and extend through clefts but never invade the lung parenchyma. Notably, they do not appear to connect to any internal patterning elements such as airways or vessels. This phenomenon of ectopic bundles of SMCs at the pleural surface has been previously reported in patients with TBX4 syndrome (Fig. 6D; (19)) but their clinical consequence is poorly understood.

**Figure 6:**
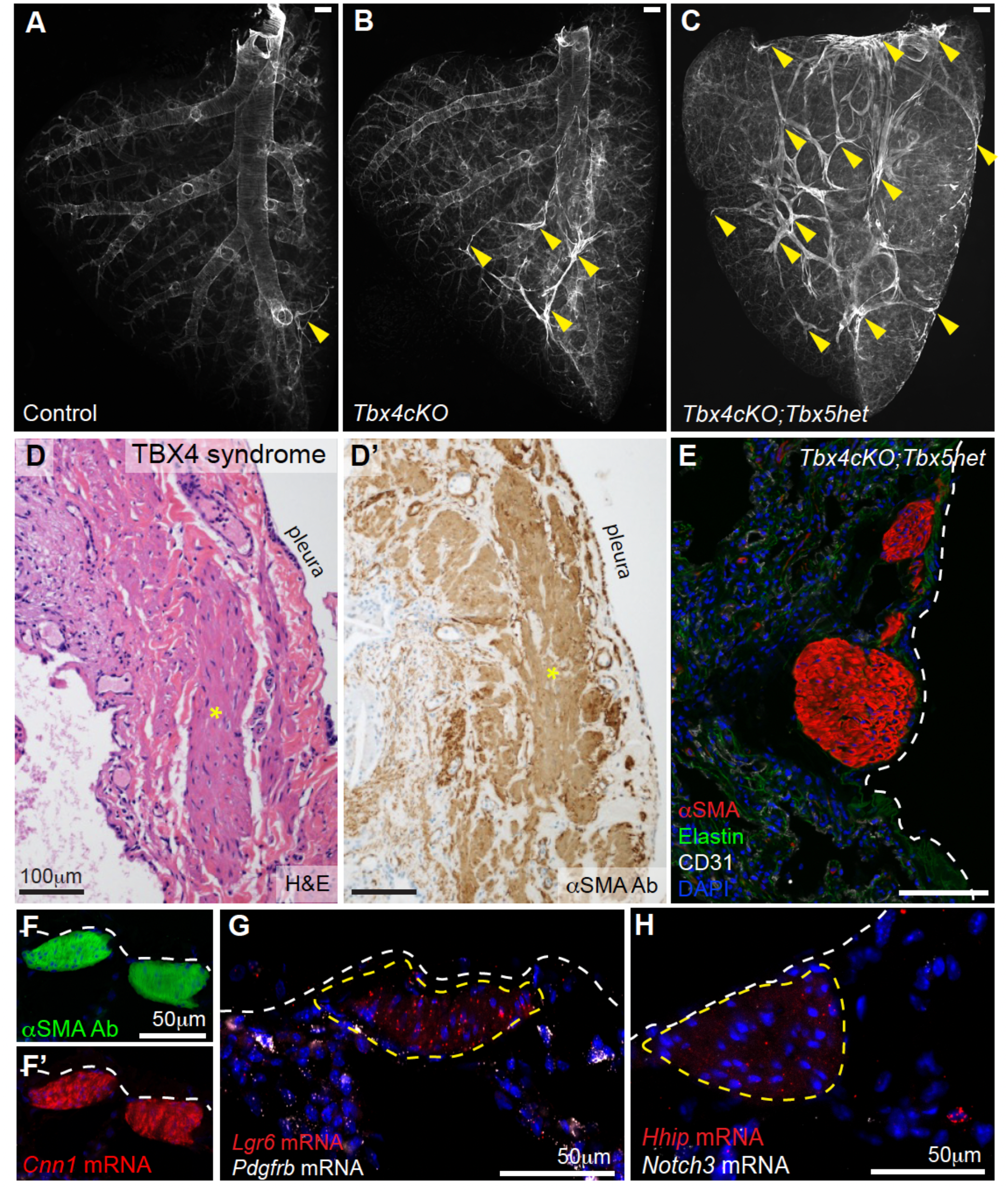
Loss of T-box genes leads to expansion of ectopic subpleural smooth muscle bands. **A**-**C**, Ventral views of right caudal lobes from control, *Tbx4cKO* and *Tbx4cKO;Tbx5het*, stained in whole mount to identify αSMA+ cells (white). Small bands of subpleural smooth muscle (arrowheads) are occasionally found in control specimens, with the size and extent of the subpleural smooth muscle banding increasing in *Tbx4cKO*, and with *Tbx4cKO;Tbx5het* animals showing extensive banding wrapping all faces of the lobe. **D&D’**, Subpleural accumulation of αSMA+ smooth muscle (asterisks) is a feature of TBX4 syndrome. **E**, Confocal micrograph of immunostained cryosection shows that mouse αSMA+ bands lie directly beneath the pleura and are composed of bundles of tightly aligned cells that lack apparent contact with internal lung structures. Molecular characterization of ectopic subpleural bands in a *Tbx4cKO;Tbx5het* mouse by fluorescent *in situ* hybridization shows αSMA+ bands express high levels of *Cnn1* (**F&F’**) and *Lgr6* (**G**), minimal levels of *Hhip* (**H**), while *Notch3* and *Pdgfrb* are undetectable, not fully consistent with any canonical lung smooth muscle identity, but intermediate between ASM and myofibroblast. Pleura marked with white dotted line, E-H; subpleural band cells marked with yellow dotted line, G&H. Scale bars 500μm, A-C; 100μm, E.

To further characterize the molecular profile of pleural bands, we performed multiplexed RNA *in situ* hybridization using markers for mesenchymal subsets identified by recent single cell profiling efforts (28), including airway and vascular SMCs, pericytes, and myofibroblasts (Fig. 6F-H). Our analysis revealed that these cells strongly express transcripts characteristic of generic SMC identity (*Cnn1*; Fig. 6F) as well as *Lgr6* (Fig. 6G), a marker of embryonic airway SMCs (25) and a lineage marker for septation myofibroblasts (41). We observed low levels of airway SMC marker *Hhip* (Fig. 6H) and negligible expression of vascular SMC and pericyte markers *Pdgfrb* and *Notch3* (Fig. 6 G&H). Collectively, this expression pattern reflects a generic αSMA-positive identity with some molecular features resembling airway SMCs and myofibroblasts, but not precisely matching any described SMC subset present in the healthy lung.

## DISCUSSION

In this study we develop a mouse model of TBX4-related lung disease that employs a lung mesenchyme-specific deletion strategy to bypass the embryonic lethality of global loss of *Tbx4*, enabling investigation of the late embryonic and adult pulmonary consequences of TBX4 syndrome *in vivo*. In this model we comprehensively characterize the changes to both the arterial network and the smooth muscle lineages of the lung that result from *Tbx4* loss of function. The model recapitulates key aspects of TBX4 syndrome, including hemodynamic evidence of PH and characteristic histopathologic findings in the pulmonary vasculature, airways, and pleura, with features worsening postnatally (9, 18-20). Given evidence of a progressive phenotype, this model offers an invaluable tool to study the mechanisms by which TBX4-related lung disease develops embryonically and worsens in postnatal life.

We provide several novel contributions to the understanding of TBX4-related lung disease. Combining tissue-specific mouse genetics, echocardiographic hemodynamic assessment, investigation of both late embryonic and adult timepoints, detailed microscopy, immunohistochemistry, and RNA *in situ* hybridization, we establish translational connections between this model and human TBX4 syndrome. Our three-dimensional approach reveals complex structural alterations that would be missed in traditional two-dimensional analyses, including elaboration of the arterial branching network, abnormal distal muscularization of both arteries and airways, and the presence of ectopic subpleural smooth muscle bands, all features of TBX4 syndrome not previously described in a mouse model. Furthermore, this study represents the first comprehensive assessment of late embryonic artery and airway SMC distribution in homozygous *Tbx4* mutants, providing crucial insights into the developmental origins and postnatal disease progression in TBX4 syndrome.

The temporal aspects of our findings are particularly relevant to understanding human disease progression (19). We observe early evidence of subtle but significant shortening of the distance between muscularized arteries and airways to the pleural edge in E18.5 embryos. This, combined with reduced lobe volumes observed in *Tbx4cKO* embryos at E18.5, prior to formation of alveoli, could suggest underdevelopment of distal airspaces, consistent with the modest reduction in branching tips observed in global *Tbx4* conditional null mice at E13.5 (21). This observation initially appears to align with Maldonado-Velez *et al*’s recent findings in a similar lung specific *Tbx4* conditional knockout model, which demonstrated enlarged alveoli, decreased vessel number, and increased vessel wall thickness in adult mice, attributing PH to disrupted alveologenesis and vascular simplification (42).

However, our analysis reveals a more complex picture. Rather than vascular simplification, we observe an elaborated arterial bed with aberrant muscularization. Specifically, our findings reveal that *Tbx4cKO* mice have smaller, more numerous muscularized arteries with thicker medial layers, and extension of muscularization further along the arterial tree that becomes even more pronounced with additional T-box gene loss. Together, these observations suggest a mechanism of increased SMC recruitment to arteries, thickening the medial layer, and aberrantly extending SMC coverage to the tiniest arterioles abutting the capillary network. These findings are consistent with a true intrinsic PAH phenotype rather than PH due to an underdeveloped vascular bed.

The smooth muscle wrapping airways is also markedly abnormal, with widened spacing of smooth muscle rings and extension of sparse αSMA+ bands far into the lung parenchyma, reaching toward the pleural surface and running with the arteries. Experiments to delineate the border between the conducting and respiratory epithelium in these altered airways, and identify how these changes relate to the alveolarization defect described by Maldonado-Velez *et al*, is needed (42). The physiologic consequences of the airway-associated smooth muscle changes we observe are unknown though muscularized conducting airways close to the pleura have been described in TBX4 syndrome (19).

We observed C-shaped αSMA-positive cells consistent with myofibroblasts in the parenchyma of *Tbx4cKO* lungs. These myofibroblasts were absent in controls, and more prominent in *Tbx4cKO;Tbx5het*. Their distribution resembles the pattern of airway-associated αSMA-positive ‘septation myofibroblasts’ present during alveologenesis (postnatal day 4-30). These cells originate as tightly parallel Lgr6-positive SMC bundles wrapping embryonic airways, which then separate and rearrange to enable epithelial budding and the formation of alveoli (41). Septation myofibroblasts downregulate αSMA expression by P30, and are not visualized by αSMA staining of adult controls (41). The distribution of αSMA-positive myofibroblasts in adult *Tbx4cKO* and *Tbx4cKO;Tbx5het* mimics that of septation myofibroblasts present during early postnatal life. This suggests *Tbx4* and *Tbx5* may be critical for the appropriate suppression of myofibroblast smooth muscle gene expression at the completion of alveologenesis.

A further example of abnormal αSMA+ cell distribution is the appearance of elaborate subpleural bands in the *Tbx4cKO* and *Tbx4cKO;Tbx5het* lungs. The molecular characterization of these subpleural structures reveals they do not correspond to any SMC subtype normally present in healthy lung tissue, though they exhibit more myofibroblast and airway smooth muscle-like features than markers typical of vascular SMCs and pericytes (28). These bands represent another manifestation of a broad dysregulation of mesenchymal cell fate and a propensity among mesenchymal cells to differentiate to an SMC identity with loss of Tbx gene function. The precise mechanisms triggering subpleural band development, the molecular signals controlling their expansion, the precise identity of their cell of origin, and their clinical impact remain undefined.

The muscularization of the arteries, dysregulation of airway-associated smooth muscle, persistence of αSMA+ myofibroblasts, and elaboration of subpleural bands each represent examples of increased smooth muscle acquisition or retention with the loss of *Tbx4*. These four cases point to *Tbx4* as a master regulator of mesenchymal differentiation whose loss leads to overabundance of αSMA+ cells in multiple compartments. The mechanism by which this SMC phenotype is acquired appears to differ between compartments, and future studies interrogating shared and divergent downstream signaling networks in each distinct mesenchymal subset in this model are needed.

This study has several limitations as a model of TBX4 syndrome. First, by design, we are investigating a pure lung phenotype that lacks compounding effects from alterations to other organs, particularly the allantois and placenta, and therefore we have necessarily lost any comorbid contributions which may worsen disease in humans (43). Further, this study was not powered to assess sex-based differences, though, contrary to other causes of PH, there is no evidence for female predominance in TBX4-related PH (44). In addition, while the combination of *TBX4* and *TBX5* mutation is not observed in human TBX4 syndrome, a large portion of TBX4 syndrome is secondary to a microdeletion that includes both *TBX4* and its homolog *TBX2*. Like *TBX4*, both *TBX2* and *TBX5* are expressed in the lung mesenchyme (23, 24, 27, 28) and some level of functional redundancy between the homologs is expected. However, small case series have not shown a worsening phenotype with increased T-box loss (19). Aligning with this, while loss of *Tbx5* led to significant SMC patterning changes, we saw no statistically significant worsening of PH hemodynamics or medial thickening in *Tbx4cKO;Tbx5het* compared to *Tbx4cKO*. Due to their proximity on Chromosome 11, dual deletion of *Tbx4* and *Tbx2* in mouse models is not feasible with Cre-lox technology.

In conclusion, our findings reveal Tbx4 functions as a key suppressor of smooth muscle differentiation within the pulmonary mesenchyme, with its loss driving excessive smooth muscle acquisition across multiple compartments in the lung. This model reveals a phenotype that mirrors the clinical pulmonary spectrum of TBX4 syndrome in humans, with embryonic origins that progressively worsen postnatally (18, 19). Three-dimensional whole mount analysis and high-resolution spatial quantitation uncover complex structural alterations that would remain hidden in traditional two-dimensional histologic approaches. This spatial precision allows us to definitively demonstrate that *Tbx4* loss results not in vascular simplification, but rather in an abnormally elaborated and over-muscularized arterial tree. Most significantly, this data demonstrates that TBX4-related pulmonary hypertension represents not merely a fixed developmental defect but rather a progressive postnatal pathophysiologic process characterized by ongoing mesenchymal dysregulation. This mechanistic insight suggests a potential therapeutic window when pharmacologic inhibition of aberrant smooth muscle differentiation and proliferation pathways could be targeted to attenuate disease progression.

## References

1. Simonneau G, Montani D, Celermajer DS, Denton CP, Gatzoulis MA, Krowka M, et al. Haemodynamic definitions and updated clinical classification of pulmonary hypertension. European Respiratory Journal. 2019 -01-24;53(1).

2. Guignabert C, Aman J, Bonnet S, Dorfmüller P, Olschewski AJ, Pullamsetti S, et al. Pathology and pathobiology of pulmonary hypertension: current insights and future directions. European Respiratory Journal. 2024 -10-31;64(4).

3. Valapour M, Lehr CJ, Schladt DP, Smith JM, Swanner K, Weibel CJ, et al. OPTN/SRTR 2022 Annual Data Report: Lung. American Journal of Transplantation. 2024 /02/01;24(2).

4. Humbert M, Guignabert C, Bonnet S, Dorfmüller P, Klinger JR, Nicolls MR, et al. Pathology and pathobiology of pulmonary hypertension: state of the art and research perspectives. European Respiratory Journal. 2019;53(1):1801887.

5. Welch CL, Austin ED, and Chung WK. Genes that drive the pathobiology of pediatric pulmonary arterial hypertension. Pediatric Pulmonology. 2021 /03/01;56(3).

6. Welch CL, and Chung WK. Genetics and Other Omics in Pediatric Pulmonary Arterial Hypertension. Chest. 2020 Jan 30;157(5).

7. Zhu N, Swietlik EM, Welch CL, Pauciulo MW, Hagen JJ, Zhou X, et al. Rare variant analysis of 4241 pulmonary arterial hypertension cases from an international consortium implicates FBLN2, PDGFD, and rare de novo variants in PAH. Genome Medicine. 2021;13(1).

8. Prapa M, Lago-Docampo M, Swietlik EM, Montani D, Eyries M, Humbert M, et al. First Genotype–Phenotype Study in TBX4 Syndrome: Gain-of-Function Mutations Causative for Lung Disease. American Journal of Respiratory and Critical Care Medicine. 2022 -12-15;206(12).

9. Thoré P, Girerd B, Jaïs X, Savale L, Ghigna M-R, Eyries M, et al. Phenotype and outcome of pulmonary arterial hypertension patients carrying a TBX4 mutation. European Respiratory Journal. 2020 -05-14;55(5).

10. Eyries M, Montani D, Nadaud S, Girerd B, Levy M, Bourdin A, et al. Widening the landscape of heritable pulmonary hypertension mutations in paediatric and adult cases. European Respiratory Journal. 2019 -03-14;53(3).

11. Chapman DL, Garvey N, Hancock S, Alexiou M, Agulnik SI, Gibson-Brown JJ, et al. Expression of the T-box family genes, Tbx1–Tbx5, during early mouse development. Developmental Dynamics. 1996 /08/01;206(4).

12. Naiche LA, Arora R, Kania A, Lewandoski M, and Papaioannou VE. Identity and fate of Tbx4-expressing cells reveal developmental cell fate decisions in the allantois, limb, and external genitalia. Developmental Dynamics. 2011 /10/01;240(10).

13. He P, Lim K, Sun D, Pett JP, Jeng Q, Polanski K, et al. A human fetal lung cell atlas uncovers proximal-distal gradients of differentiation and key regulators of epithelial fates. Cell. 2022 /12/08;185(25).

14. Zhang B, He P, Lawrence JEG, Wang S, Tuck E, Williams BA, et al. A human embryonic limb cell atlas resolved in space and time. Nature 2023 635:8039. 2023 -12-06;635(8039).

15. Bongers EMHF, Duijf PHG, Beersum SEMv, Schoots J, Kampen Av, Burckhardt A, et al. Mutations in the Human TBX4 Gene Cause Small Patella Syndrome. The American Journal of Human Genetics. 2004 /06/01;74(6).

16. Haarman MG, Kerstjens-Frederikse WS, and Berger RMF. TBX4 variants and pulmonary diseases: getting out of the ‘Box’. Current Opinion in Pulmonary Medicine. May 2020;26(3).

17. Karolak JA, Welch CL, Mosimann C, Bzdęga K, West JD, Montani D, et al. Molecular Function and Contribution of TBX4 in Development and Disease. American Journal of Respiratory and Critical Care Medicine. 2022 Nov 11;207(7).

18. Karolak JA, Vincent M, Deutsch G, Gambin T, Cogné B, Pichon O, et al. Complex Compound Inheritance of Lethal Lung Developmental Disorders Due to Disruption of the TBX-FGF Pathway. The American Journal of Human Genetics. 2019 /02/07;104(2).

19. Galambos C, Mullen MP, Shieh JT, Schwerk N, Kielt MJ, Ullmann N, et al. Phenotype characterisation of TBX4 mutation and deletion carriers with neonatal and paediatric pulmonary hypertension. European Respiratory Journal. 2019 -08-22;54(2).

20. Doughty ES, Norvik C, Levin A, Bodmer J, Tran-Lundmark K, Abman SH, et al. Long-Term Effect of TBX4 Germline Mutation on Pulmonary Clinico-Histopathologic Phenotype. Pediatric and Developmental Pathology. 2024 -1;27(1).

21. Arora R, Metzger RJ, and Papaioannou VE. Multiple Roles and Interactions of Tbx4 and Tbx5 in Development of the Respiratory System. PLOS Genetics. Aug 2, 2012;8(8).

22. Guo M, Morley MP, Jiang C, Wu Y, Li G, Du Y, et al. Guided construction of single cell reference for human and mouse lung. Nature Communications 2023 14:1. 2023 -07-29;14(1).

23. Shirazi SP, Negretti NM, Jetter CS, Sharkey AL, Garg S, Kapp ME, et al. Bronchopulmonary Dysplasia with Pulmonary Hypertension Associates with Loss of Semaphorin Signaling and Functional Decrease in FOXF1 Expression. bioRxiv. 2024 -09-01.

24. Zanini F, Che X, Suresh NE, Knutsen C, Klavina P, Xie Y, et al. Hyperoxia prevents the dynamic neonatal increases in lung mesenchymal cell diversity. Scientific Reports 2024 14:1. 2024 -01-23;14(1).

25. Kumar ME, Bogard PE, Espinoza FH, Menke DB, Kingsley DM, and Krasnow MA. Defining a mesenchymal progenitor niche at single-cell resolution. Science. 2014 -11-14;346(6211).

26. Zhang W, Menke DB, Jiang M, Chen H, Warburton D, Turcatel G, et al. Spatial-temporal targeting of lung-specific mesenchyme by a Tbx4enhancer. BMC Biology 2013 11:1. 2013 -11-13;11(1).

27. Negretti NM, Plosa EJ, Benjamin JT, Schuler BA, Habermann AC, Jetter CS, et al. A single-cell atlas of mouse lung development. Development. 2021 /12/15;148(24).

28. Travaglini KJ, Nabhan AN, Penland L, Sinha R, Gillich A, Sit RV, et al. A molecular cell atlas of the human lung from single-cell RNA sequencing. Nature 2020 587:7835. 2020 -11-18;587(7835).

29. Naiche LA, and Papaioannou VE. Loss of Tbx4 blocks hindlimb development and affects vascularization and fusion of the allantois. Development. 2003 /06/15;130(12).

30. Rallis C, Bruneau BG, Del Buono J, Seidman CE, Seidman JG, Nissim S, et al. Tbx5 is required for forelimb bud formation and continued outgrowth. Development. 2003 /06/15;130(12).

31. Wabel EA, Krieger-Burke T, and Watts SW. Vascular chemerin from PVAT contributes to norepinephrine and serotonin-induced vasoconstriction and vascular stiffness in a sex-dependent manner. American Journal of Physiology-Heart and Circulatory Physiology. 2024 Dec 05.

32. Steffes LC, Froistad AA, Andruska A, Boehm M, Mcglynn M, Zhang F, et al. A Notch3-Marked Subpopulation of Vascular Smooth Muscle Cells Is the Cell of Origin for Occlusive Pulmonary Vascular Lesions. Circulation. 2020;142(16):1545–61.

33. Shen Z, Lu Z, Chhatbar PY, O’Herron P, Kara P, Shen Z, et al. An artery-specific fluorescent dye for studying neurovascular coupling. Nature Methods 2012 9:3. 2012 -01-22;9(3).

34. Arora R, Fries A, Oelerich K, Marchuk K, Sabeur K, Giudice LC, et al. Insights from imaging the implanting embryo and the uterine environment in three dimensions. Development. 2016 /12/15;143(24).

35. Madhavan MK, DeMayo FJ, Lydon JP, Joshi NR, Fazleabas AT, and Arora R. Aberrant uterine folding in mice disrupts implantation chamber formation and alignment of embryo-uterine axes. Development. 2022 /06/01;149(11).

36. Brittain E, Penner NL, West J, and Hemnes A. Echocardiographic Assessment of the Right Heart in Mice. Journal of Visualized Experiments (JoVE). 2013 -11-27(81).

37. Ma Z, Mao L, and Rajagopal S. Hemodynamic Characterization of Rodent Models of Pulmonary Arterial Hypertension. J Vis Exp. 2016 (110).

38. Potus F, Martin AY, Snetsinger B, and Archer SL. Biventricular Assessment of Cardiac Function and Pressure-Volume Loops by Closed-Chest Catheterization in Mice. J Vis Exp. 2020 (160).

39. Metzger RJ, Klein OD, Martin GR, Krasnow MA, Metzger RJ, Klein OD, et al. The branching programme of mouse lung development. Nature 2008 453:7196. 2008 -05-07;453(7196).

40. Wagenvoort CA, and Wagenvoort N. Pathology of Pulmonary Hypertension. John Wiley & Sons; 1977.

41. Gillich A, Julien KRS, Brownfield DG, Travaglini KJ, Metzger RJ, and Krasnow MA. Alveoli form directly by budding led by a single epithelial cell. bioRxiv. 2021 -12-26.

42. Maldonado-Velez G, Mickler EA, Cook TG, and Aldred MA. Loss of Tbx4 Affects Postnatal Lung Development and Predisposes to Pulmonary Hypertension. bioRxiv. 2024 -09-22.

43. Mestan KK, Check J, Minturn L, Yallapragada S, Farrow KN, Liu X, et al. Placental Pathologic Changes of Maternal Vascular Underperfusion in Bronchopulmonary Dysplasia and Pulmonary Hypertension. Placenta. 2014 May 20;35(8).

44. Zhu N, Gonzaga-Jauregui C, Welch CL, Ma L, Qi H, King AK, et al. Exome Sequencing in Children With Pulmonary Arterial Hypertension Demonstrates Differences Compared With Adults. Circulation: Genomic and Precision Medicine. 2018 April;11(4).

