## Supplemental Data for "Excessive Postnatal Smooth Muscle Differentiation in a Lung Specific Model of *TBX4*-related Pulmonary Hypertension"

**Supplemental Figure 1. Characterization of Tbx4 deletion in embryonic and postnatal mice. A**, Semi-quantitative PCR was performed using three primers targeting the Tbx4 conditional (“floxed”) and excised conditional (“null”) alleles. The conditional allele was effectively excised in Cre-positive lungs, as indicated by the marked decrease in intensity of the conditional band and presence of the null band. Non-mesenchymal cells carry floxed alleles without Cre expression; therefore, the presence of a conditional band is expected in both *Tbx4cKO* and conditional samples. As a control, the null band was generated using only the two primers targeting the null allele in a *Tbx4cKO* sample. **B**, No significant difference in body weight was found between control, *Tbx4cKO*, and *Tbx4cKO;Tbx5he*t adult mice at the time of harvest. **C**, PCR primer sequences (Tbx4-F, Tbx4-M, and Tbx4-R) used for genotyping and verification of conditional *Tbx4* allele excision.

**Supplemental Figure 2. Left heart function is not significantly affected by loss of T-box alleles.** No significant difference in cardiac output (CO), stroke volume (SV) or ejection fraction (EF) between control, *Tbx4cKO* and *Tbx4cKO;Tbx5het* mice.

Supplementary Figure 1

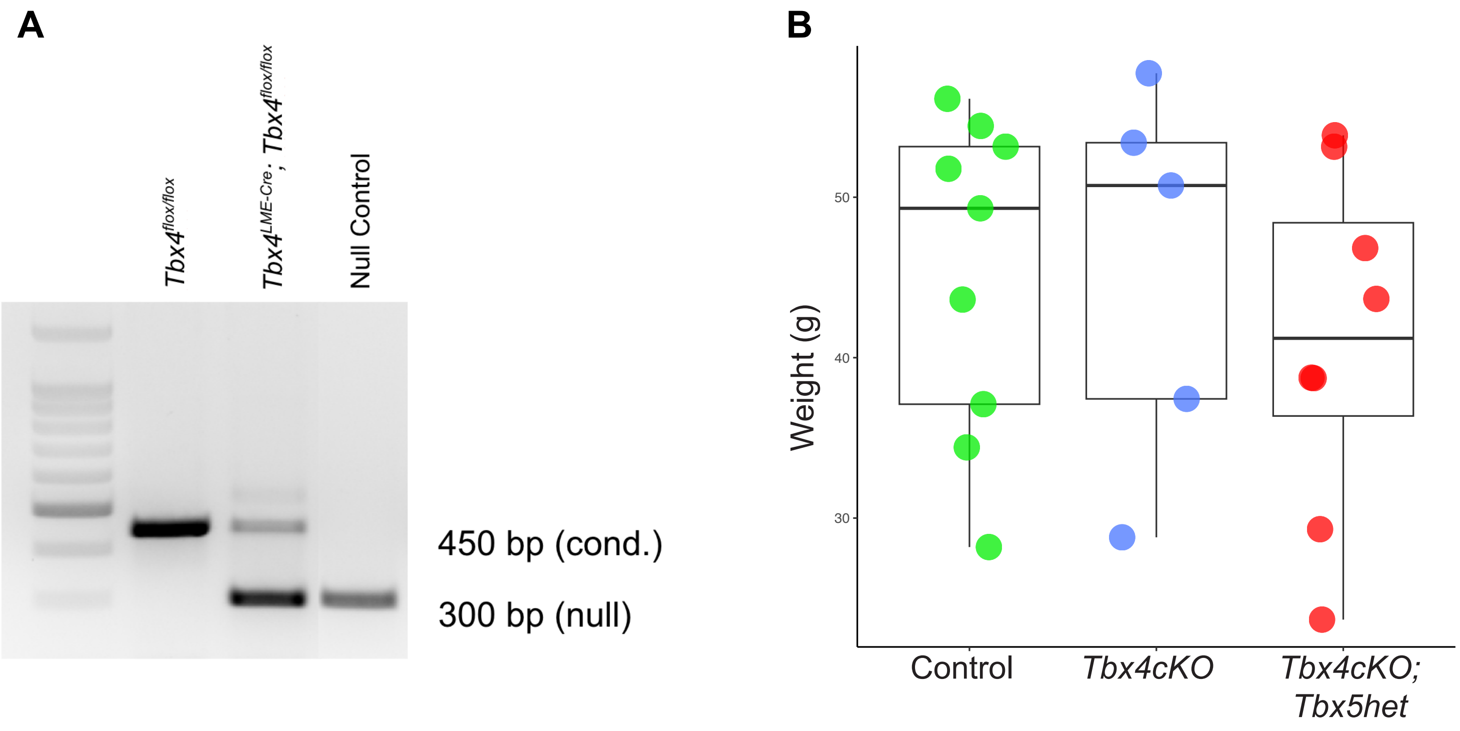


**C** PCR primers used to genotype and assess conditional allele excision.

| **Primer** | **Sequence** |
|---|---|
| Tbx4 Conditional Allele Excision | |
| Tbx4-F | GCAGATGACCATCGCTACAA |
| Tbx4-M | TCATCTAGGCTTCACAGCC |
| Tbx4-R | GCCAGGCAGAGAAACAAAAG |

Supplementary Figure 2

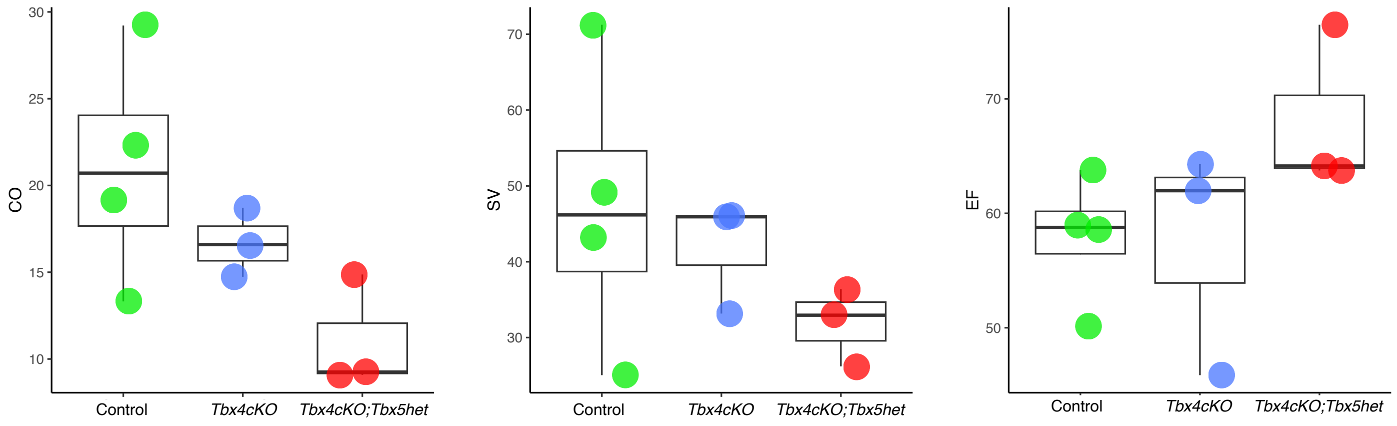
